# High-performance xenograft read-sorting for DNA methylation analysis

**DOI:** 10.64898/2026.10.06.756952

**Authors:** Stefan Altendorfer, Jens Zentgraf, Johanna Elena Schmitz, Christoph Plass, Clarissa Gerhäuser, Sven Rahmann

## Abstract

Xenograft sorting tools separate sequenced DNA reads by species. We have enhanced the computational method and tool Xengsort to enable efficient epigenetic profiling of patient-derived xenograft models; it now supports cytosine-converted whole genome or reduced-representation data. Benchmarking Xengsort against several established methods (ICRG, bamcmp, XenofilteR) revealed consistently high accuracy across tools, but Xengsort outperformed the other tools in computational efficiency due to its alignment-free design.

**Graphical Abstract:** 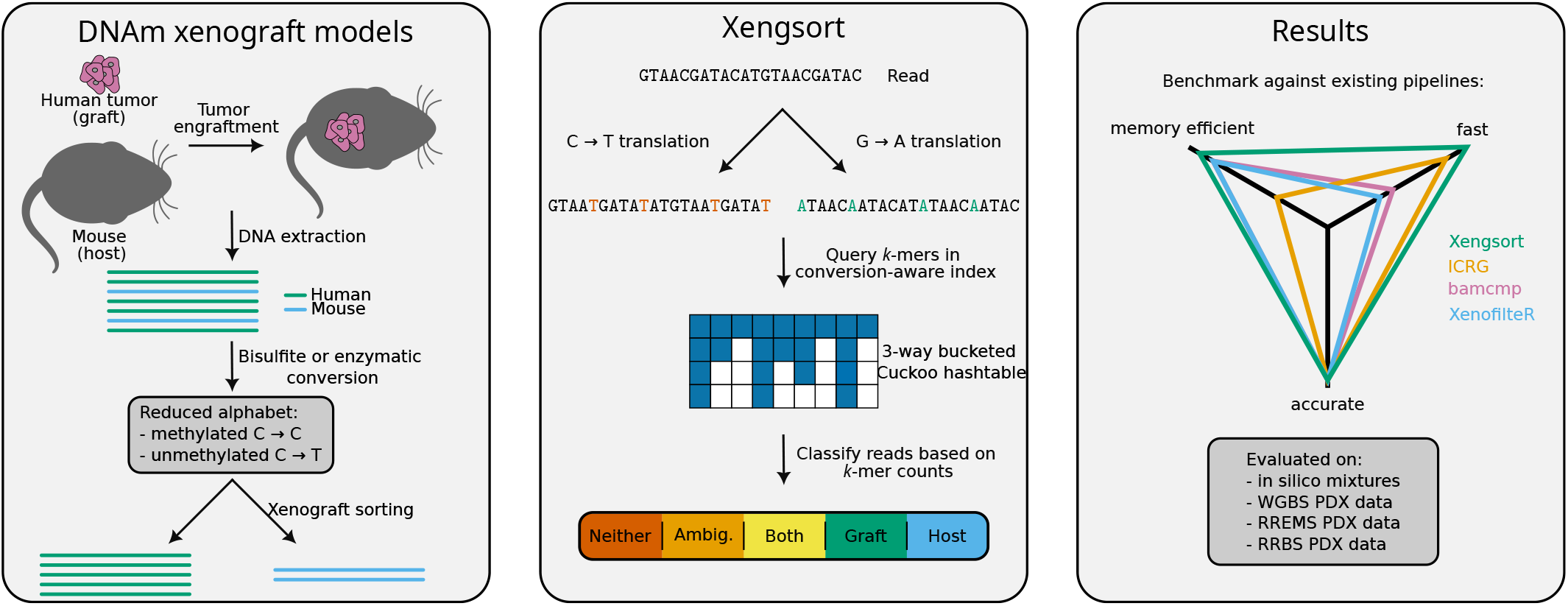

## Introduction

Xenografts are created by transplanting cells, tissues, or entire organs from one species into another. Patient-derived xenograft (PDX) mouse models are widely used in cancer research to investigate tumor progression, therapy response, and mechanisms of drug resistance in a physiological in vivo context that preserves most of the tumor’s cellular architecture and heterogeneity. (Blanchard et al., 2025). Following successful engraftment and tumor growth, the xenografts are harvested and undergo next-generation sequencing. However, the analysis of xenograft samples is complicated by the presence of host contamination originating from the mouse microenvironment (Qian et al., 2022). Recently, epigenetic profiling has become important for understanding how tumors adapt to treatment and evolve under therapeutic pressure (Romero-Garcia et al., 2020). DNA methylation (DNAm) is a well-studied regulatory process that plays a central role in tumor initiation, lineage specification, and therapy resistance (Smith et al., 2025; Marine et al., 2020). Chemical or enzymatic DNA conversion methods enable genome-wide characterization of DNAm by converting unmethylated cytosines to uracil (sequenced as thymine), while methylated cytosines remain protected from conversion. This cytosine conversion substantially reduces sequence complexity and alters nucleotide composition, posing challenges for accurate read alignment and species classification (Lin et al., 2025), as conversion-aware aligners (Pedersen et al., 2014) map a substantial amount of mouse reads onto the human reference genome (Extended Data Fig. 1).

Various read-sorting methods have been developed to separate human from (non-human) mouse reads in genomic and transcriptomic sequencing datasets. The most recent benchmarking winner (Bhandari et al., 2025), Xengsort (Zentgraf and Rahmann, 2021), is a pre-alignment-based method for classifying xenograft-derived sequence reads into five classes: host, graft, both, neither, and ambiguous. Instead of expensively computing pairwise alignments, it uses host/graft/both-labeled short subsequences of length *k*, indexed in a highly optimized hash table built from both reference genomes. An alternative read-sorting approach involves sorting reads during alignment by using an in silico combined reference genome (ICRG), which allows the aligner to determine whether a read primarily aligns to a human or a mouse chromosome. Post-alignment read-sorting tools, such as bamcmp (Khandelwal et al., 2017) and XenofilteR (Kluin et al., 2018), operate on two BAM files generated from separate alignments against the human and the mouse reference genomes. These methods then compare the mapping scores of each read/read-pair across these two alignments to identify its species of origin.

While these existing read-sorting approaches work well for conventional sequencing data (Bhandari et al., 2025), they have mostly not yet been evaluated for cytosine-converted reads. Some tools, such as Xengsort, could previously not be applied to DNA methylation data since cytosine conversion alters the sequence identity required for *k*-mer matching.

To address this problem, we here have adapted Xengsort to handle converted reads for index building and classification. For a systematic benchmark, we applied alignment-based (ICRG), and post-alignment-based methods (bamcmp, XenofilteR) to DNAm data to assess their performance across various in-silico human–mouse mixtures and on multiple real PDX datasets.

## Materials and Methods

### Basic definitions

Given a sequence *s* over the 4-nucleotide DNA alphabet Σ_DNA_ = {A, C, G, T} with length *n* = |*s*|, a *k*-mer of *s* is a substring of length *k*. There are (*n* − *k* + 1) *k*-mers in *s*, of which some may be equal. To compute the reverse complement *rc*(*x*) of a sequence or a *k*-mer *x*, we reverse it and replace each nucleotide with its complement (A with T; C with G).

A *k*-mer can be efficiently represented using 2-bit encoding as follows: A ↦ (00)_2_ = 0_10_, C ↦ (01)_2_ = 1, G → (10)_2_ = 2, T → (11)_2_ = 3. The integer encoding *enc*(*x*) of a *k*-mer is then the value of the concatenated 2*k*-bit representation, e.g., ATC ↦ (001101)_2_ = 13. Since a *k*-mer *x* and its reverse complement *rc*(*x*) represent the same molecule, we use the maximum of their encodings as a representative for both, called its canonical code *cc*(*x*) = max(*enc*(*x*), *enc*(*rc*(*x*))). This allows for treating both DNA strands simultaneously with an efficient numerical encoding.

### The xenograft sorting problem

Given a short sequence *s* (a sequenced read) from a sample that contains mixed reads from a host (mouse) and a graft species (human), the problem of xenograft-sorting is to classify this read as originating from either *host* or *graft*. Three additional classes allow for a more fine-grained classification: *both*, which contains all reads that may originate from the host or the graft, e.g., if a read originates from a region that is similar or identical in both reference genomes; *neither* if a read is neither similar to the host reference nor the graft reference; *ambiguous* if one part of the read uniquely maps to the host reference and another part uniquely maps to the graft reference. Common reasons for a *neither* classification are common contaminations, such as reads from the Epstein–Barr virus, or spike-in for sequencing, such as reads from the PhiX phage.

### Xengsort

Xengsort is an alignment-free method to classify reads from xenograft samples using a *k*-mer index, mapping each *k*-mer to species information (occurs in host, graft, both) (Zentgraf and Rahmann, 2021).

#### Indexing

For a given host and graft species, the first step is to build the index once, which can then be used to process all samples. For indexing, all *k*-mers are extracted from the host and graft reference genomes and inserted into a multi-way bucketed Cuckoo hash table, which is a fast and memory-efficient key-value data structure. Together with each *k*-mer, it stores the information in which reference genome(s) the *k*-mer occurs. A *k*-mer is unique if it occurs in only one of the species (otherwise it is non-unique). We further partition the unique *k*-mers into strongly unique and weakly unique ones. A *k*-mer is strongly unique if no similar *k*-mer (at Hamming distance 1, i.e., one nucleotide is changed) occurs in the other species; otherwise, it is weakly unique. Three bits are used to encode the value of a *k*-mer (1 bit for contained in host; 1 bit for contained in graft; 1 bit for strong/weak distinction). During classification, we trust strongly unique *k*-mers more than weakly unique *k*-mers, as they are more robust towards variants or sequencing errors.

For DNA methylation sequencing of xenograft samples, we extended the index of Xengsort to contain all *k*-mers that can be obtained after bisulfite or enzymatic conversion. For this, we translate each *k*-mer of the reference genomes into a *k*-mer where all Cs are replaced with Ts and, for the reverse complementary strand, into a *k*-mer where all Gs are replaced with As (Fig. 1a).

**Figure 1.**
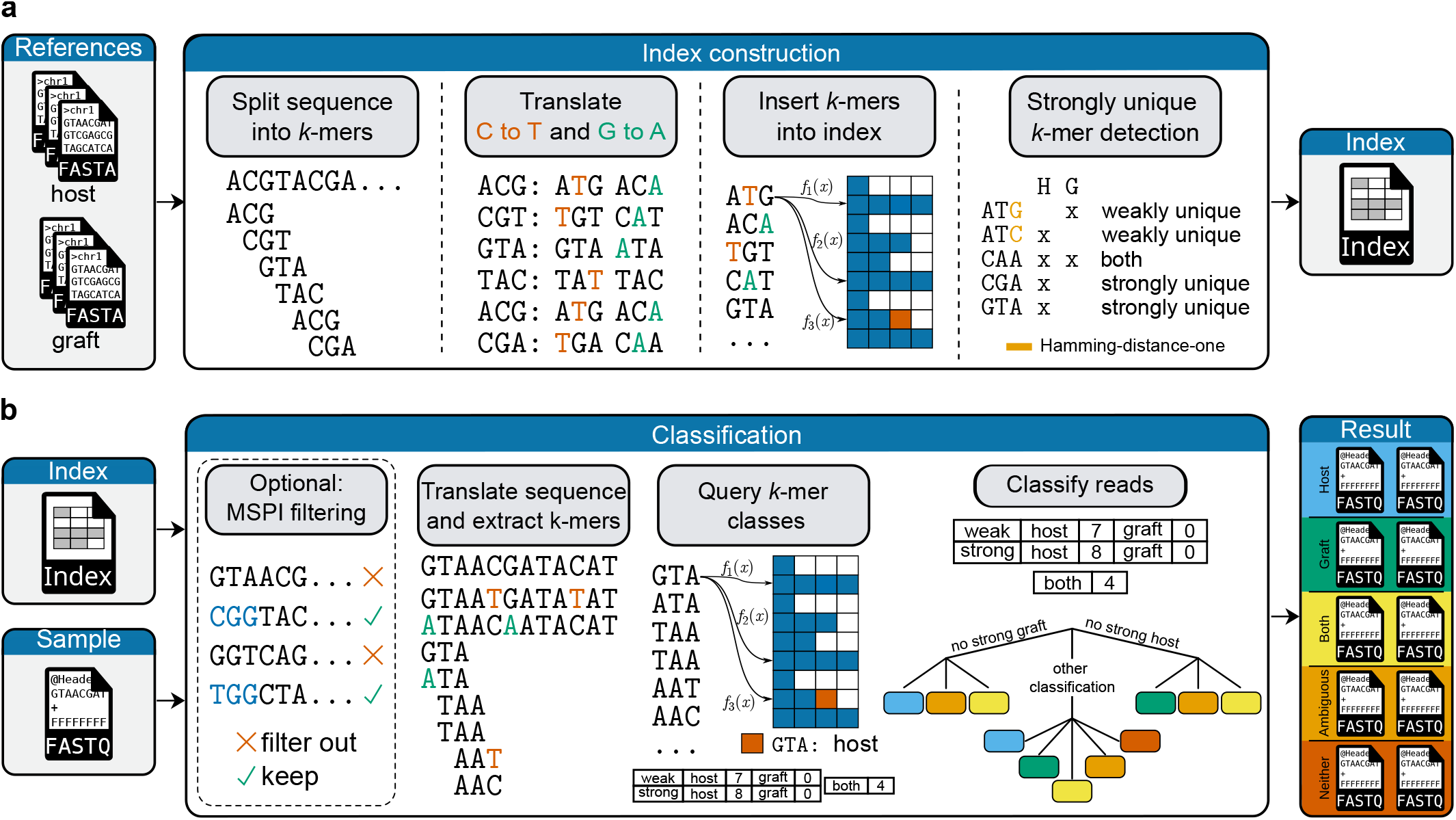
Xengsort index construction (a) and classification step (b) with alphabet conversion for DNA methylation analysis a. Mouse (host) and human (graft) reference genomes are split into *k*-mers, and two variants of each *k*-mer (C→T and G→A translated) are inserted into a fast and memory-efficient 3-way bucketed Cuckoo hash table index. Xengsort classifies each *k*-mer as strongly unique (only in human or mouse, no similar *k*-mer at Hamming distance one in the other species), weakly unique (only in human or mouse, but with similar *k*-mer in the other species), and non-unique (occurs in both human and mouse). b. For species assignment of any single read, the species information (human, mouse, both, neither) of each translation of each of the read’s *k*-mers is obtained from the index. The assignment is made by a classification tree based on *k*-mer type statistics. For reduced-representation sequencing data, Xengsort provides the option to filter reads without the restriction site prior to read classification.

The index is built once and stored on disk and can be loaded into memory. Alternatively, it may reside in shared memory and be used by several read classification tasks simultaneously.

#### Read classification from FASTQ files

For read classification, we process each read twice, under both C→T and G→A translation. We extract all *k*-mers and query them against the respective index (Fig. 1b). Based on these counts, we first determine the translation type of the read. For most reads, most *k*-mers will be stored in the index (host, graft, or both) for one translation, while for the other translation, most *k*-mers will not be found. Thus, if the number of *k*-mers found for one conversion is at least twice as high as for the other conversion, we consider only the counts for this conversion to classify the read into one of the following categories: host, graft, both, ambiguous, or neither. Otherwise, we use the combined counts to classify the read. For single-read sequencing, the counts are classified using the same rules as described for the original Xengsort (Zentgraf and Rahmann, 2021). For paired-end sequencing, the first and second reads are classified independently using the same rules as used for single-read sequencing, and the final classification is decided based on both classifications according to Table 1.

**Table 1.**
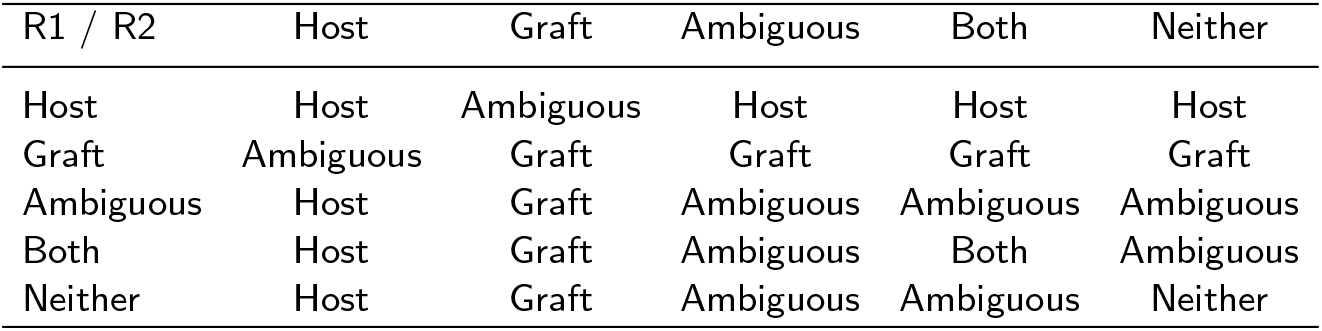
Classification of paired-end reads based on the classification results for read 1 (R1) and read 2 (R2). The table is symmetric.

| R1 / R2 | Host | Graft | Ambiguous | Both | Neither |
| --- | --- | --- | --- | --- | --- |
| Host | Host | Ambiguous | Host | Host | Host |
| Graft | Ambiguous | Graft | Graft | Graft | Graft |
| Ambiguous | Host | Graft | Ambiguous | Ambiguous | Ambiguous |
| Both | Host | Graft | Ambiguous | Both | Ambiguous |
| Neither | Host | Graft | Ambiguous | Ambiguous | Neither |

After classification, the read is written to the respective class-specific FASTQ output file, such that the final output of Xengsort is five FASTQ files (2× five for paired-end data) containing the reads of the respective classes.

#### Restriction to MspI sites

When processing Reduced-Representation Bisulfite-Sequencing (RRBS) or EM-seq data, only reads that originate from an MspI start site need to be processed. During classification, we offer an optional filtering step for these. Before the read classification process is started (which includes looking up all *k*-mers of the read), it is first checked whether the read starts with an MspI site pattern (CGG or TGG recognition sequence). If this is the case, the read is classified. Otherwise, the read is discarded and stored in an additional output file.

### Data preparation

#### In silico read mixture

Whole-genome bisulfite sequencing (WGBS) datasets for in-silico benchmarking were downloaded from the NCBI Sequence Read Archive (SRA) (Leinonen et al., 2011) using the corresponding GEO accessions listed in Supplementary Table 1. Raw FASTQ files were retrieved using the SRA Toolkit (v3.2.1) (Leinonen et al., 2011).

In silico human–mouse mixtures were generated from adapter-trimmed paired-end FASTQ files using seqkit (v2.13.0) (Shen et al., 2024). Mixtures were produced at two sequencing depths (10M and 50M read pairs), and nine human-mouse ratios with six replicates per condition. For each downsampling replicate, human and mouse reads were subsampled independently using identical random seeds for read 1 and read 2 to preserve pairing relationships. Subsampled reads were concatenated to produce mixed FASTQ files. FastQC reports were generated for quality control, and paired-end reads were adapter-trimmed and quality-filtered using Trim Galore (v0.6.10) (Martin, 2011) with identical parameters across all datasets. Reads were conservatively trimmed with a Phred quality threshold of 20, and fixed clipping was applied to remove 10 bp from both the 5^*′*^ and 3^*′*^ ends of each mate (R1 and R2). To retain ground-truth species origin after merging, read identifiers were prefixed with “human” or “mouse” prior to concatenation.

All mixtures were processed using the nf-core/methylseq pipeline (v4.1.0-gdb60bfe) (Ewels et al., 2025) implemented in Nextflow (v25.10.0) (Di Tommaso et al., 2017) with Singularity containerization (Kurtzer et al., 2017) to ensure computational reproducibility. Alignment was performed using the bwa-meth aligner (v0.2.9) (Pedersen et al., 2014) with the BWA-MEM2 backend enabled (Vasimuddin et al., 2019-05). Adapter trimming was skipped in the pipeline (`--skip trimming`) because reads had been uniformly preprocessed prior to mixture generation. The `--fasta` parameter was set to the appropriate reference build for each species.

Post-alignment processing, including sorting and indexing, was performed using samtools (v1.21) (Danecek et al., 2021). Default pipeline parameters were used unless otherwise specified.

#### PDX models

To apply the tools to real DNA methylation data, WGBS data for breast cancer PDX models (GSE237768) (Achinger-Kawecka et al., 2024), and RREMS and RRBS data for breast cancer PDX, patient-derived xenograft organoids (PDXO), and patient tumor samples (GSE152202, GSE186747) (Guillen et al., 2022) were downloaded from the NCBI SRA database (Supplementary Table 2).

Adapter trimming and quality filtering were performed using Trim Galore (v0.6.10) (Martin, 2011) with dataset-specific parameters following the recommendations of the original authors. For WGBS data (GSE237768), paired-end reads were trimmed with a minimum length threshold of 40 bp, with 6 bp clipped from the 3^*′*^ end of R2 and 6 bp from the 3^*′*^ end of R1. For RRBS data (GSE152202), the `--rrbs` mode was applied, while paired-end RREMS data (GSE186747) was processed with the additional `--paired` flag.

Bisulfite sequencing alignments were performed using the nf-core/methylseq pipeline (v4.1.0)(Ewels et al., 2020) with RefSeq reference genomes and parameters identical to those described in the “Parameter settings for read-sorting tools” section.

An additional optional pre-filtering step was applied to the RRBS and RREMS datasets to assess the efficacy of restriction enzyme digestion.

All four read-sorting strategies (Xengsort, ICRG, bamcmp, and XenofilteR) were tested to enable enable comparison in real biological PDX datasets. XenofilteR’s MM-threshold was kept at the default value of four for RRBS and RREMS, as recommended by the tutorial for datasets with short read lengths (50 bp).

### Parameter settings for read-sorting tools

Four read-sorting strategies were evaluated: bamcmp (Khandelwal et al., 2017), XenofilteR (Kluin et al., 2018), Xengsort (Zentgraf and Rahmann, 2021), and an in silico combined reference genome (ICRG) approach (Callari et al., 2018).

For alignment-based approaches (bamcmp and XenofilteR), reads were aligned separately to the human (GRCh38.p14) and mouse (GRCm38) NCBI RefSeq reference genomes using the nf-core/methylseq workflow described above.

Bamcmp (v2.2) species assignment was performed with name-based comparison (`-n`) and match scoring mode (`-s match`). The reads are partitioned into four categories based on alignment quality across the two reference genomes: human-only, human-better, mouse-only, and mouse-better. For performance evaluation, reads classified as human-only or human-better were considered human, whereas reads classified as mouse-only or mouse-better were considered non-human.

XenofilteR was run in R (v1.6) using the `XenofilteR()` function with default parameters, except that the MM-threshold (maximal edit distance between read and reference genome) was evaluated at 4 and 8 for 100 bp and 150 bp paired-end data. A threshold of 8 performed better, with reduced human read loss. The method only retains reads classified as human. Since the library size and mixture proportions were known for the in silico benchmarking, misclassified human and non-human reads could be calculated for later accuracy evaluations.

For the ICRG strategy, competitive alignment was performed against a concatenated human–mouse reference genome constructed by merging GRCh38.p14-genomic and GRCm38 assemblies. Mouse chromosome identifiers were prefixed with “m.” to enable unambiguous species identification following alignment. Reads were assigned to species based on their primary alignment target within the combined reference or otherwise classified as unmapped. Unmapped reads or reads mapped to mouse chromosomes were considered as non-human for accuracy score evaluation.

Xengsort (v2.2.1) was executed in classification mode using a pre-built bisulfite-converted human–mouse *k*-mer index. Reads were processed directly from trimmed paired-end FASTQ files without prior alignment. For accuracy evaluation, reads labelled as graft were considered human, whereas reads labelled as host, ambiguous, both, or neither were considered non-human for in silico benchmarking.

### Performance evaluation

#### Measuring runtime and memory usage

For illustration, wall-clock runtime and peak memory (`peak rss`) usage were exemplarily evaluated using mixtures containing 60% human reads at a sequencing depth of 50 million paired-end reads across all four dataset pairs. Total runtime was defined as the sum of read-sorting and execution of the nf-core/methylseq workflow, including alignment and BAM processing steps.

Alignment was performed using the `--use mem2` and `--fasta` RefSeq references parameters (`BWAMETH ALIGN`), while BAM processing included sorting and indexing (`SAMTOOLS SORT`, `SAMTOOLS INDEX ALIGNMENTS`). For bamcmp and XenofilteR, both independent alignments to the human and mouse reference genomes were summed in runtime.

For the ICRG strategy, runtime and maximum memory were determined solely by the alignment against the concatenated human–mouse reference and subsequent BAM processing.

Although Xengsort is alignment-free, to ensure comparability of downstream analysis times, reads classified as human were additionally aligned to the human reference genome using the same alignment workflow described above. The reported runtime and memory usage for Xengsort, therefore, includes both classification and subsequent human-only alignment.

All alignment steps were performed using nf-core/methylseq with default process configurations (12 CPUs for `BWAMETH ALIGN`). read-sorting tools were run with 8 threads (Xengsort, bamcmp), except XenofilteR, which does not support multi-threading and was therefore run single-threaded. Three technical replicates were executed per condition. Reported runtimes correspond to per-replicate wall-clock execution time.

#### Accuracy score

Sorting performance was evaluated relative to the known ground-truth species origin encoded in the read identifiers. Because the primary objective in xenograft analysis is the accurate recovery of human reads, the evaluation focused on high retention of human reads and low contamination rate.

Reads were categorized as follows. True positives (TP): human reads correctly classified as human; false negatives (FN): human reads incorrectly classified as non-human; false positives (FP): non-human reads incorrectly classified as human; true negatives (TN): non-human reads correctly classified as non-human.

Human read loss was defined as the fraction of true human reads incorrectly classified as non-human:]

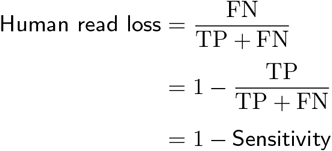

Contamination rate was defined as the fraction of true non-human reads incorrectly classified as human:

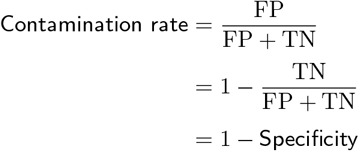

To summarize tool performance across metrics, a weighted accuracy score was computed:

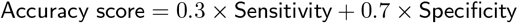

This formulation assigns greater weight to contamination, reflecting the practical importance of minimizing host-derived reads in downstream tumor methylation analyses. Lower human read loss and contamination rate indicate better overall accuracy. Weights were defined a priori and applied uniformly across all tools, mixture ratios, and sequencing depths.

#### Efficiency score

To summarize computational efficiency, the runtime and peak memory usage were each min–max normalized to the interval [0, 1] across tools and inverted so that higher values denote better runtime or memory efficiency:

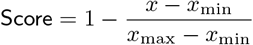

Then, the overall efficiency score was defined as the equally weighted mean of runtime score and memory usage score, each on a [0, 1] scale as described above.

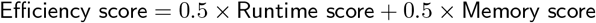

#### Total score

To obtain an overall ranking, accuracy and computational efficiency were combined into a total score:

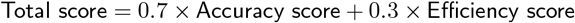

Weights were defined a priori and applied uniformly across all tools. Higher scores indicate better overall performance.

## Results

### Optimal *k*-mer length for Xengsort

In the adapted Xengsort, we nearly double the number of *k*-mers in the index because it contains both C→T and G→A translated *k*-mers (Fig. 2A). This also results in a higher number of weakly unique *k*-mers and *k*-mers that occur in both species. As the *k*-mer length increases, the *k*-mers become more specific, and the proportion of strongly unique *k*-mers increases in both the normal and cytosine-converted indices. Consequently, the proportion of reads classified as ambiguous decreases with increasing *k*-mer length (Fig. 2B). We therefore determined that a *k*-mer length of 29, rather than 25 as in the original implementation, was optimal for cytosine-converted index building.

**Figure 2.**
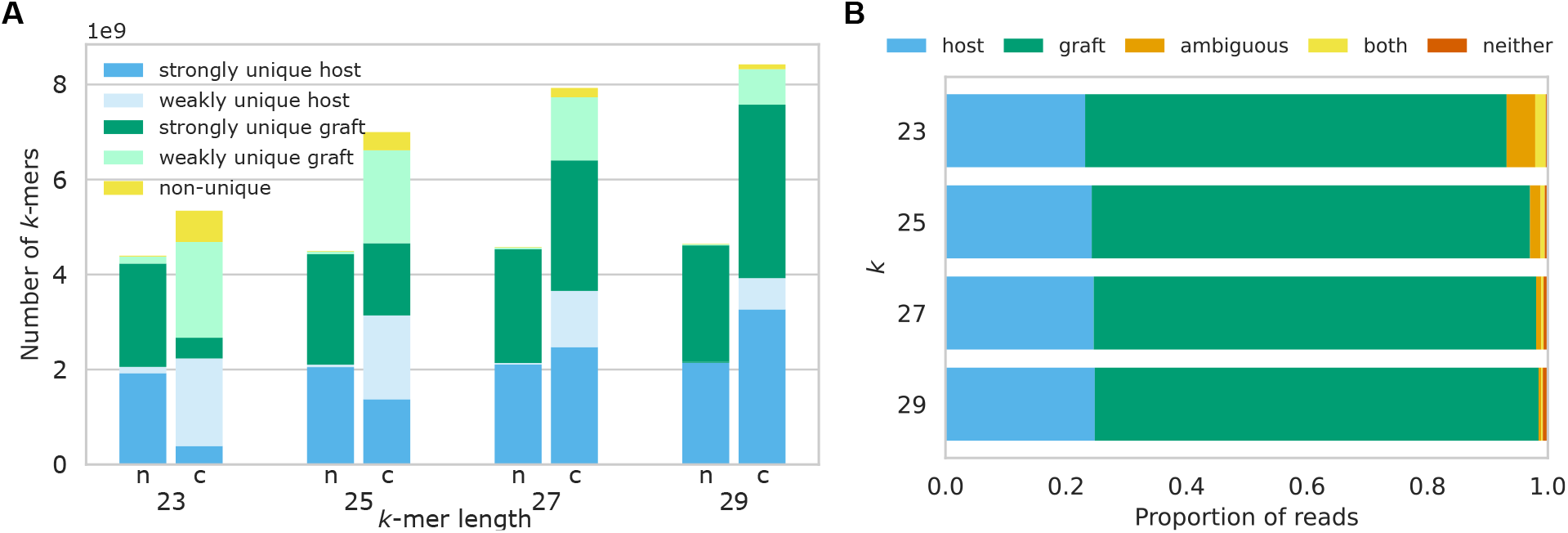
Xengsort index and classification for different *k*-mer lengths. Absolute numbers of *k*-mers for different values of *k* in the combined human-mouse index, according to *k*-mer type: strongly unique host/graft (only in human or mouse, no Hamming-distance-one *k*-mer in the other species), weakly unique in host/graft (only in human or mouse, but a Hamming-distance-one *k*-mer exists in the other species), and non-unique (in human and mouse). Numbers are given for both the standard index (n) and the cytosine-converted index (c; contains C*→*T and G*→*A translated *k*-mers). Proportion of reads classified as host, graft, both, ambiguous, and neither for different *k*-mer lengths (HCI-005-Decitabine-1 dataset, accession GSM7648678).

### Xengsort achieves competitive accuracy with substantially lower computational cost

To benchmark the performance of Xengsort, we compared its performance against other read-sorting methods (ICRG, bamcmp, and XenofilteR) for the application on cytosine-converted reads. ICRG has previously been used for representation-based methylation sequencing only, but not in a genome-wide context. The two post-alignment tools bamcmp and XenofilteR have not been applied to DNAm before, but are technically applicable to the conversion-aware wrapper of bwa-mem (Pedersen et al., 2014).

To assess the sorting performance across these tools, we created a series of in silico human–mouse mixtures (see *Materials and Methods*) to enable direct quantification of read-sorting accuracy. Four human–mouse dataset pairs were selected to represent biologically and technically distinct sample types. Mixture 1 combined a human breast cancer cell line dataset (Carmona et al., 2014) and primary mouse bone marrow hematopoietic stem cells (Sun et al., 2014), both sequenced at 100 bp read length. Mixtures 2–4 were derived from human and mouse pancreatic organoid datasets (Wang et al., 2024) sequenced at 150 bp. Two total sequencing depths were evaluated: 50 million and 10 million paired-end reads per mixture (corresponding to 100 million and 20 million single-end reads, respectively). For each dataset pair, reads were randomly subsampled at predefined human-content mixture ratios ranging from 10% to 90% (in increments of 10%). For each mixture ratio and sequencing depth, six independent replicates were generated using distinct random seeds (0–5).

To evaluate read-sorting accuracy, we applied the following metrics: Human read loss denotes the fraction of reads erroneously discarded, while contamination rate depicts the fraction of wrongly retained non-human reads after sorting. These metrics were combined into a weighted accuracy score.

When we scored the human read loss against the different mixture datasets, we observed similar trends across the 150 bp mixtures (Mix 2–4). All tools exhibited human read losses below 1.5%, except for XenofilteR, which was too stringent and discarded ∼7% of human reads (Fig. 4A). Mix 1 was a slight outlier due to its 100 bp sequencing characteristics, leading to increased human read losses (e.g., ∼3.3% and ∼2.1% for Xengsort and bamcmp, respectively). In WGBS datasets, however, losses below 5% are unlikely to affect downstream analyses due to retained coverage. Additionally, performances were unaffected by sequencing depth and human content within the mixtures.

**Figure 3.**
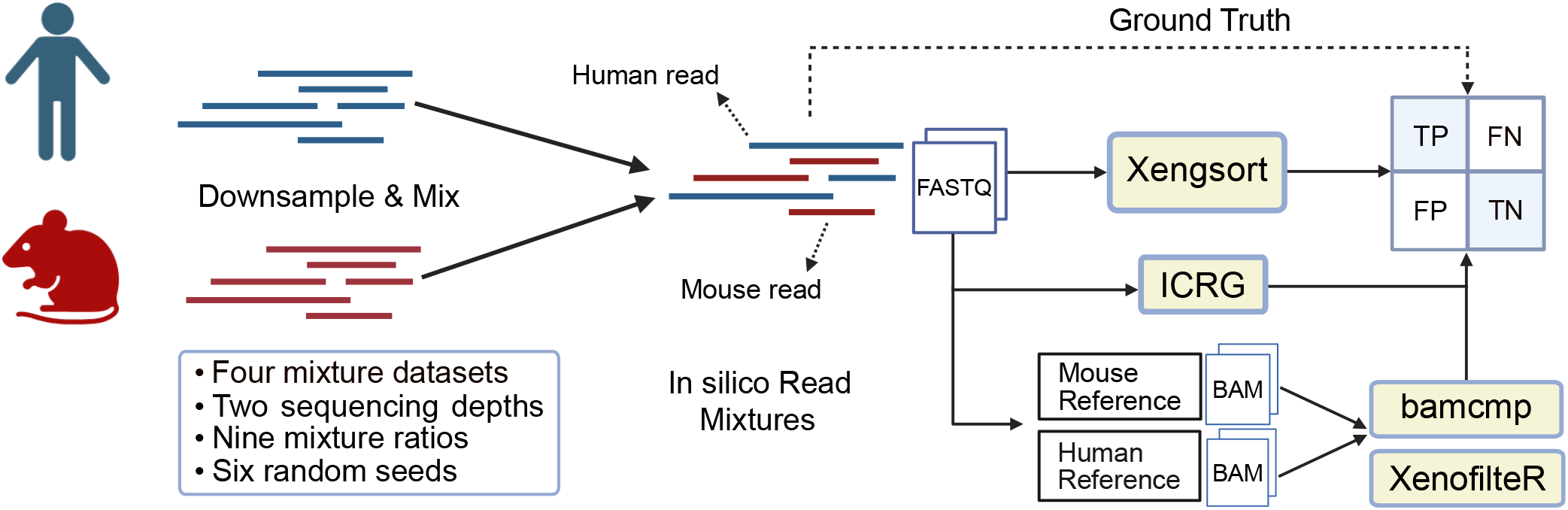
Benchmarking and validating read-sorting strategies for cytosine-converted sequencing data. Generation of in-silico read mixtures to assess the accuracy of Xengsort, ICRG, bamcmp and XenofilteR.

**Figure 4.**
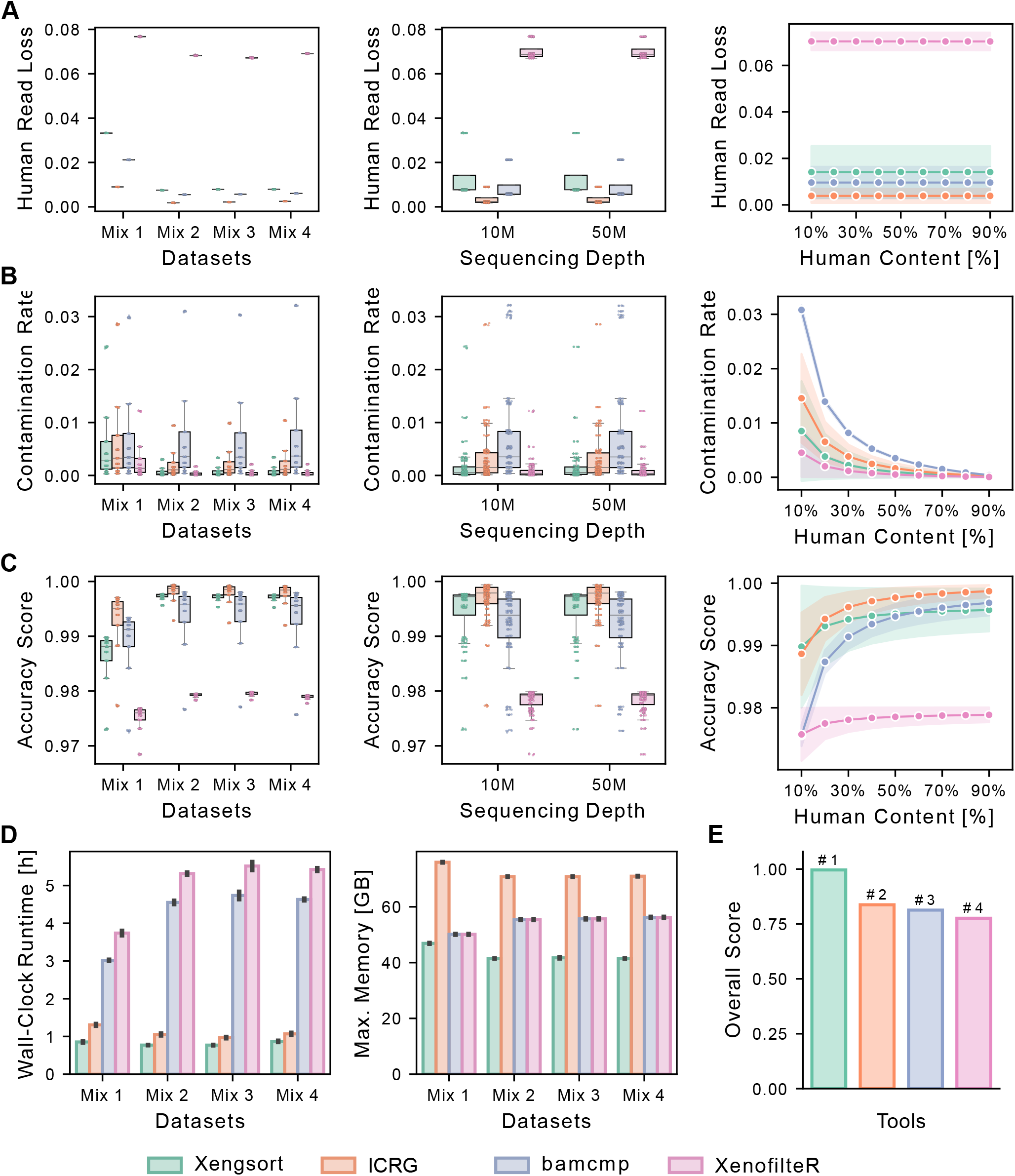
Read-sorting accuracy and computational efficiency of read-sorting workflows across in-silico benchmarking conditions. Human read loss (A), contamination rate (B), and composite accuracy score (C) for each tool, shown as a function of mixture dataset, sequencing depth, and human content. Shading around the line plots indicates standard deviations. Boxplots show the distribution of the grouped read-sorting outcomes: the box spans the interquartile range (Q1-Q3), with the median marked as a line. Lines and whiskers extend to 1.5× the interquartile range. (D) Wall-clock runtime [h] and maximal memory usage [GB] for the full read-sorting workflow (including sorting and alignment), measured across three replicates of in silico mixtures 1-4 at 60% human fraction. For Xengsort, human-classified reads were aligned against GRCh38. For Bamcmp and XenofilteR, reads were aligned separately against GRCm38 and GRCh38, and the runtimes were summed. For ICRG, reads were aligned against a concatenated reference genome. Error bars define the standard deviation across replicates. (E) The final ranking (overall score) integrates the accuracy score (C) and efficiency (D) into an overall score (see *Materials and Methods*).

We observed a slight increase in contamination rate for Mix 1, whereas Mix 2–4 behave similarly (Fig. 4B). Sequencing depth did not alter the results. Residual contamination, however, decreased with increasing human content for every tool, reflecting the smaller pool of mouse reads available to escape filtering. While Xengsort and XenofilteR showed mostly contamination rates below 1%, ICRG and bamcmp exhibited higher rates of contamination, peaking at 1.5% and 3%, respectively, when the human content was low.

By combining human read loss and contamination rate into a weighted composite accuracy score, Xengsort, ICRG and bamcmp achieved values of above 99%, emphasizing their comparable classification accuracy under all evaluated conditions (Fig. 4C). XenofilteR lacked closely behind (∼97%) because of its higher human read loss.

In contrast, efficiency and scalability differed markedly. Xengsort achieved the fastest runtime (below 1 hour) and lowest memory usage (∼43 GB) by performing classification at the FASTQ level, thereby aligning only human-classified reads (Fig. 4D, Extended Data Fig. 2). ICRG showed short runtimes (∼1.1 h) but had the highest memory usage (∼72 GB) because of the large combined reference index. bamcmp and XenofilteR required the longest runtimes (∼4.2 and ∼5.0h, respectively) due to dual-genome alignments while maintaining moderate memory demands (∼54 GB).

Integrating classification accuracy and efficiency into an overall score, Xengsort achieved the highest ranking among the evaluated tools (Fig. 4E). Overall, we demonstrated that all evaluated read-sorting methods achieve highly reliable results in filtering human reads from in-silico cytosine-converted read mixtures. Xengsort, however, outperformed other methods in computational efficiency due to its pre-alignment sorting workflow.

### High performance of Xengsort on real PDX data from different conversion protocols

We further compared the read-sorting tools on multiple real breast cancer (BC) PDX DNAm datasets. We downloaded WGBS xenograft data from a study that examined the effects of epigenetic treatment on endocrine-resistant metastatic breast cancer (GSE237768) (Achinger-Kawecka et al., 2024) (Fig. 5A). The study comprises two patient-derived xenograft (PDX) models originating from pleural effusion (HCI-005) and liver (Gar15-13), which were treated with either decitabine (a DNA demethylating agent) or vehicle control. While the WGBS dataset was not sorted into human and mouse reads, the authors reported that the Gar15-13 model contained ∼15–18% non-human reads in accompanying HiC data. (Achinger-Kawecka et al., 2024). Xengsort, ICRG, and bamcmp identified similar but consistently lower proportions of 4–8% host reads in the bisulfite-converted DNA from the same model (Fig. 5B). Differences in tumor purity estimation across sequencing modalities has already been report in previous studies. (Qian et al., 2022; Haider et al., 2020) XenofilteR was not applicable (NA) to this large dataset due to R memory limitations. We determined human read content ranging from 70–83% for PDX model HCl-005, again with high concordance across tools. Overall, the amount of (mouse) contamination was primarily influenced by the inherent properties of the PDX models and was independent of decitabine treatment. Comparing computational efficiency across the read-sorting workflows, we observed a strong drop in wall-clock runtime and maximal memory usage for Xengsort (5.5h, 43GB), as compared to ICRG (8.0h, 73 GB) and bamcmp (29.1h, 49 GB) (Fig. 5C). Notably, bamcmp’s runtime was largely inflated by aligning large human-dominated PDX samples against the mouse reference, where millions of reads generate spurious seed matches that the aligner must extend before discarding them.

**Figure 5.**
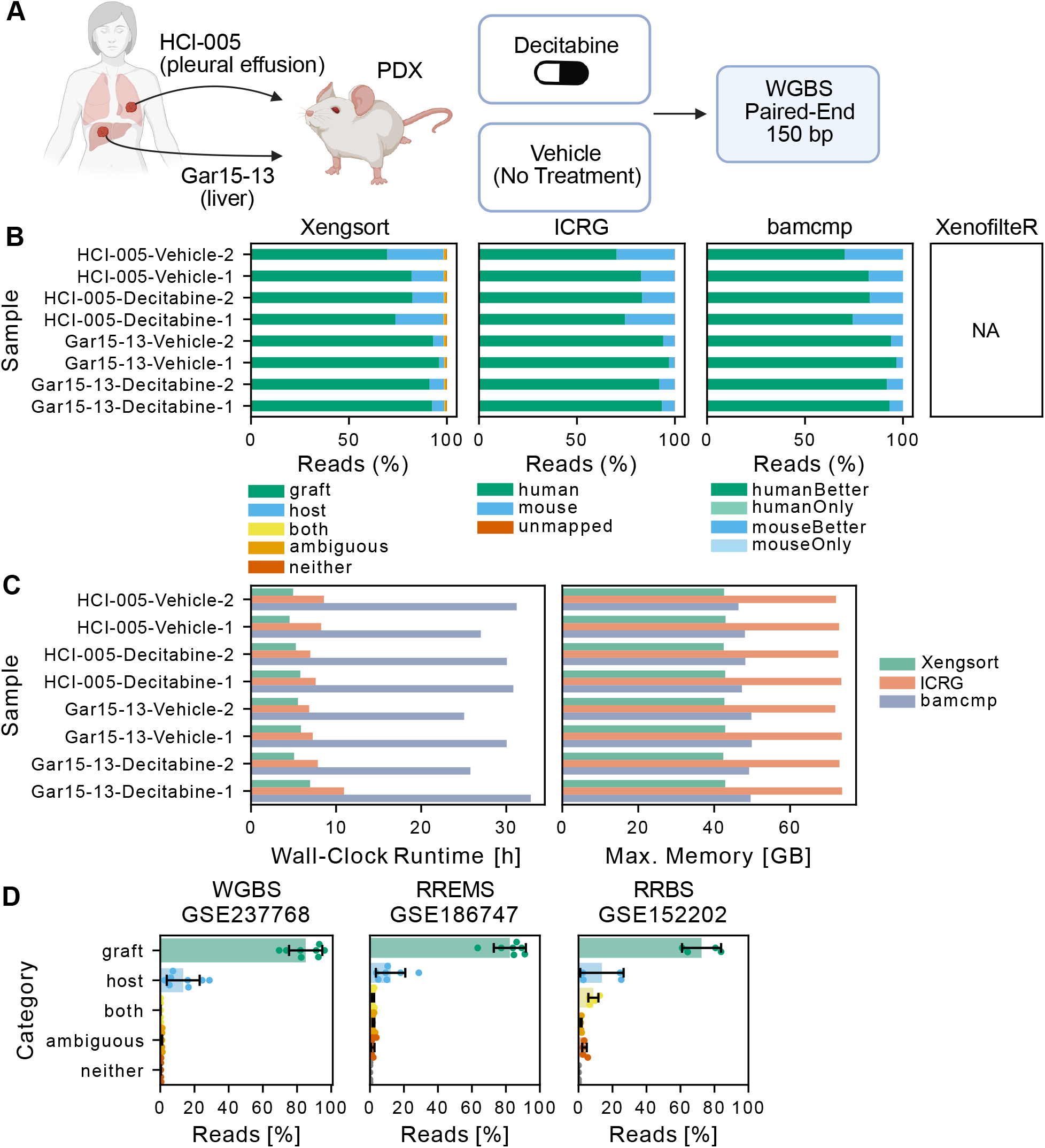
Validation of read-sorting algorithms on metastatic breast cancer PDX data A. Paired-end WGBS data for metastatic breast cancer PDX samples treated with Decitabine (DNA demethylating agent) or Vehicle (control). B. Classification of PDX reads via Xengsort, ICRG, and bamcmp. XenofilteR was not applicable (NA) due to R memory limitations. C. Sum of wall-clock runtime and maximal memory usage for the combined read-sorting and alignment step for each workflow. D. Validation of Xengsort’s applicability to public breast cancer xenograft data. Error bars indicate the standard deviation across PDX models.

We additionally applied the tools to Reduced-Representation Enzymatic Sequencing (RREMS, GSE186747) and Bisulfite Sequencing (RRBS, GSE152202) datasets that provide matched patient tumor, PDX, and patient-derived xenograft organoid (PDXO) samples from primary, recurrent, and metastatic breast cancer (Extended Data Fig. 3A, 4A). Both sequencing technologies rely on MspI (or alternative) restriction enzyme digestion to enrich CpG-rich regions, which poses a challenge for read-sorting due to evolutionary conservation (Long et al., 2013). We tested two classification modes: removal of reads lacking the MspI start site, or using all reads.

For the RREMS dataset, read-sorting tools concordantly estimated mouse contents ranging from 10–28% (Fig. 5D, Extended Data Fig. 3B) without MspI filtering. Uncertainty classes remained below 5%, aided by leveraging information from the paired read for classification. For the single-end RRBS dataset with short 50 bp reads, reads classified as “both” rose to 8%, limiting Xengsort’s *k*-mer-based approach (Fig. 5D).

MspI pre-filtering revealed significant variation in digestion efficiency across different sequencing samples. While the median of the proportion of reads lacking the MspI start site was around 9%, individual values ranged from 3–84% and 8–61% in RREMS and RRBS datasets, respectively due to a few outlier samples. Some of the PDX samples, in particular, demonstrated very low digestion efficency (e.g. HCI-003 PDX≈84%, HCI-032 PDX≈82%, HCI-027 PDX≈61%). These reads (annotated as “sites”) would predominantly have been annotated as “graft”. Efficiency and scalability analyses revealed Xengsort’s advantages in runtime over dual-alignment approaches and maximum memory usage over the concatenated reference genome. However, caution should be exercised regarding the high number of “both” classifications under single-end RRBS (Extended Data Fig. 3-4).

## Conclusion

In this study, we have established the use of several read-sorting tools for DNA methylation analysis.

Firstly, we adapted the most recent benchmarking winner, Xengsort, to account for cytosine conversion. Xengsort now generates *k*-mer indices for both reduced alphabets (A,G,{T,C} and {A,G},T,C) simultaneously, making the method conversion-aware and, at the same time, tolerant towards incomplete conversion. In the classification step, we process each read twice, under both C→T and G→A translation, querying all of its translated *k*-mers against the respective index and classifying it as host, graft, both, neither, or ambiguous. While ICRG has already been applied to reduced-representation methylation sequencing, we now demonstrate that popular post-alignment tools bamcmp and XenofilteR, can also be used with cytosine-converted data. Although most read-sorting tools achieved high precision and sensitivity under various benchmarking conditions, Xengsort showed a clear advantage in computational efficiency, which became particularly evident when applied to large-scale real xenograft datasets.

In summary, we recommend the following workflow for analysing xenograft methylation data: First, apply Xengsort to sort reads by their genome of origin. Then, perform read alignment on the graft-derived reads by using the aligner of your choice. Finally, proceed with downstream methylation processing.

### Key points

- Xengsort has been successfully adapted to handle cytosine-converted reads during *k*-mer index construction and read classification, using data from WGBS, RREMS and RRBS xenograft studies.
- Alignment-based read-sorting tools are also applicable for DNAm data with bisulfite-aware aligners.
- Xengsort, ICRG, bamcmp and XenofilteR achieve high read-sorting accuracies across varying benchmark conditions
- Xengsort’s flexible pre-alignment read-sorting workflow outperforms other methods in runtime and maximum memory usage, becoming especially remarkable in large-scale Xenograft datasets.

## Ethics declarations

LLMs (Claude, Anthropic) were used to support coding of data visualization scripts.

## Data availability

All raw whole-genome bisulfite sequencing datasets used for benchmark generation are publicly available from the NCBI Sequence Read Archive (SRA) under the accession numbers listed in Tables S1 and S2.

## Code availability

Our software, xengsort, is available via GitLab (http://gitlab.com/genomeinformatics/xengsort), PyPi (https://pypi.org/project/xengsort/), and bioconda (https://anaconda.org/channels/bioconda/packages/xengsort/overview). Code to reproduce the results of this paper is available at https://github.com/S-Altend/MethSortBench.

## Author contributions statement

J.Z., J.E.S., and S.R. designed and implemented the original and modified algorithms. S.A. conducted the performance evaluation of Xengsort for DNA methylomes and applied it to real breast cancer datasets. J.Z. and S.A. drafted the manuscript. S.A., J.Z., and J.E.S. created the figures. C.G, C.P., and S.R. edited the manuscript and provided supervision. All authors reviewed and approved the final version of the manuscript.

**Stefan Altendorfer** is currently a PhD student at the German Cancer Research Center in the Cancer Epigenomics division. His research involves bioinformatics analysis of DNA methylation in triple-negative breast cancer.

**Jens Zentgraf** is currently a PhD student at the Saarbrücken Graduate School of Computer Science in the Algorithmic Bioinformatics group of Sven Rahmann. His research interests are the development of efficient alignment-free methods for bioinformatics.

**Johanna Elena Schmitz** is currently a PhD student at the Saarbrücken Graduate School of Computer Science in the Algorithmic Bioinformatics group of Sven Rahmann. Her research interests are the development of efficient methods and statistical models for bioinformatics.

**Christoph Plass** is head of the Division of Cancer Epigenomics, and his work focuses on genetic and epigenetic alterations in leukemia.

**Clarissa Gerhäuser** is a group leader in the division Cancer Epigenomics at the German Cancer Research Center in Heidelberg. Her research focuses on epigenomic alterations associated with metastatic progression and therapy resistance in breast and prostate cancer.

**Sven Rahmann** is professor and chair of Algorithmic Bioinformatics at Saarland University. His research concerns algorithms and data structures for alignment-free biological sequence analysis and related statistical problems.

