## Extended Data Figures for "High-performance xenograft read-sorting for DNA methylation analysis"

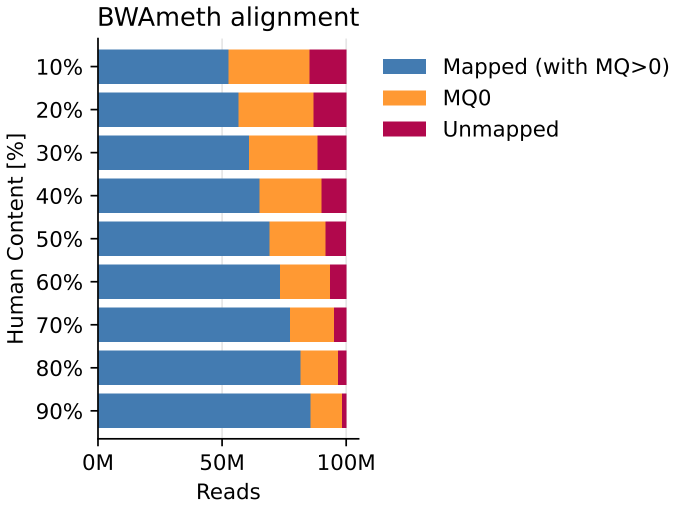


**Extended Data Fig. 1**: Mapping qualities for cytosine-converted human-mouse read mixtures against the human reference genome. The read mixtures were generated *in silico* at 50M paired-end sequencing depth from a mouse and human pancreatic cancer organoid at increasing human content from 10-90%. BWAmeth alignment reports mapping qualities of reads against GRCh38. $MQ > 0$ illustrates that the mapped reads have a degree of unique alignment confidence. $MQ0$ alignments are ambiguous, as the read maps equally well to two or more places in the reference genome. *Unmapped* reads could not be aligned to any location in the reference genome.

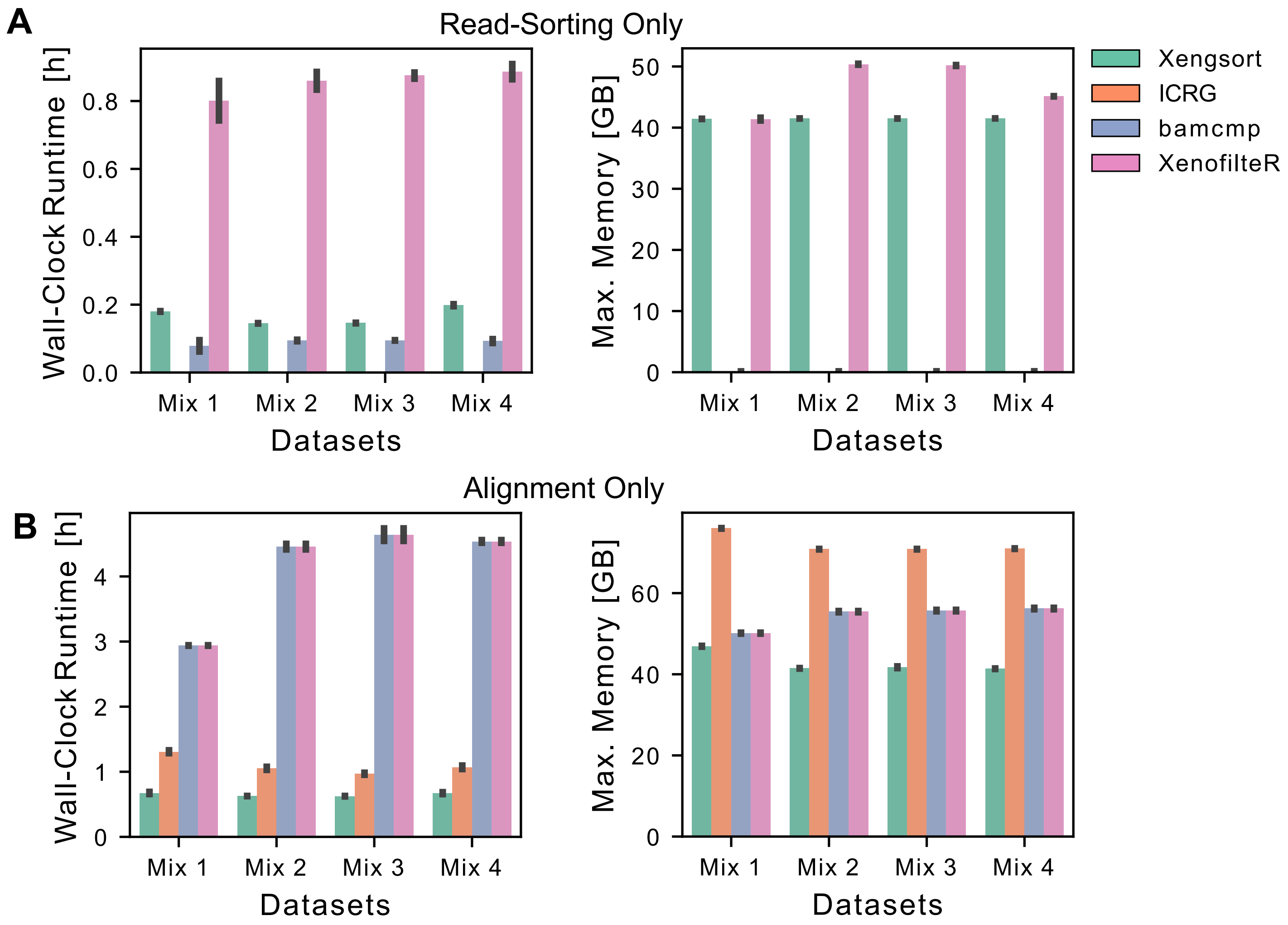


**Extended Data Fig. 2**: Efficiency and scalability analysis for read-sorting workflows in benchmarking conditions Maximum wall-clock runtime [h] and memory usage [GB] for: A. the read-sorting step and B. the alignment step, measured across three replicates of in silico mixtures 1–4 at 60% human fraction. For Xengsort, human-classified reads were aligned against GRCh38. For Bamcmp and XenofilteR, reads were aligned separately against GRCm38 and GRCh38, and the runtimes were summed. For ICRG, reads were aligned against a concatenated reference genome. Error bars define the standard deviation across replicates.

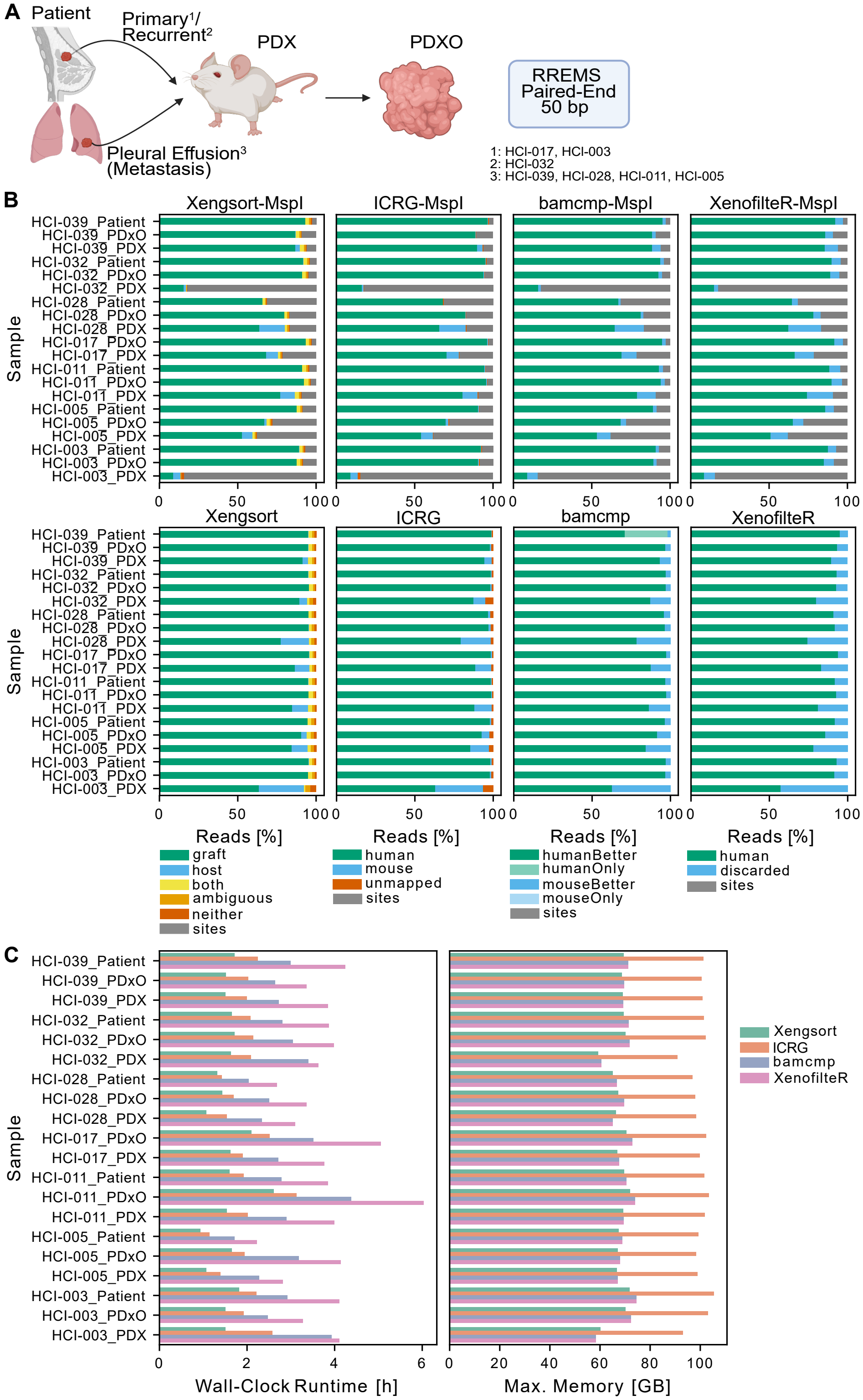


**Extended Data Fig. 3**: Validation of read-sorting algorithms on breast cancer PDX Reduced-Representation EM-Sequencing (RREMS) data (GSE186747) A. Paired-end RREMS data includes primary, recurrent, and metastatic breast cancer, PDX-derived organoid (PDXOs) and PDX samples. B. Classification of sample reads via Xengsort, ICRG, bamcmp, and XenofilteR with MspI start site prefiltering (top) or sorting of all reads (bottom). Non-MspI reads (*sites*, indicated in grey) comprise a median of 9% reads, though some samples reached up to 84%, indicating variable digestion efficiency. C. Sum of wall-clock runtime and maximal memory usage for the combined read-sorting and alignment step for each non-MspI-prefiltering workflow.

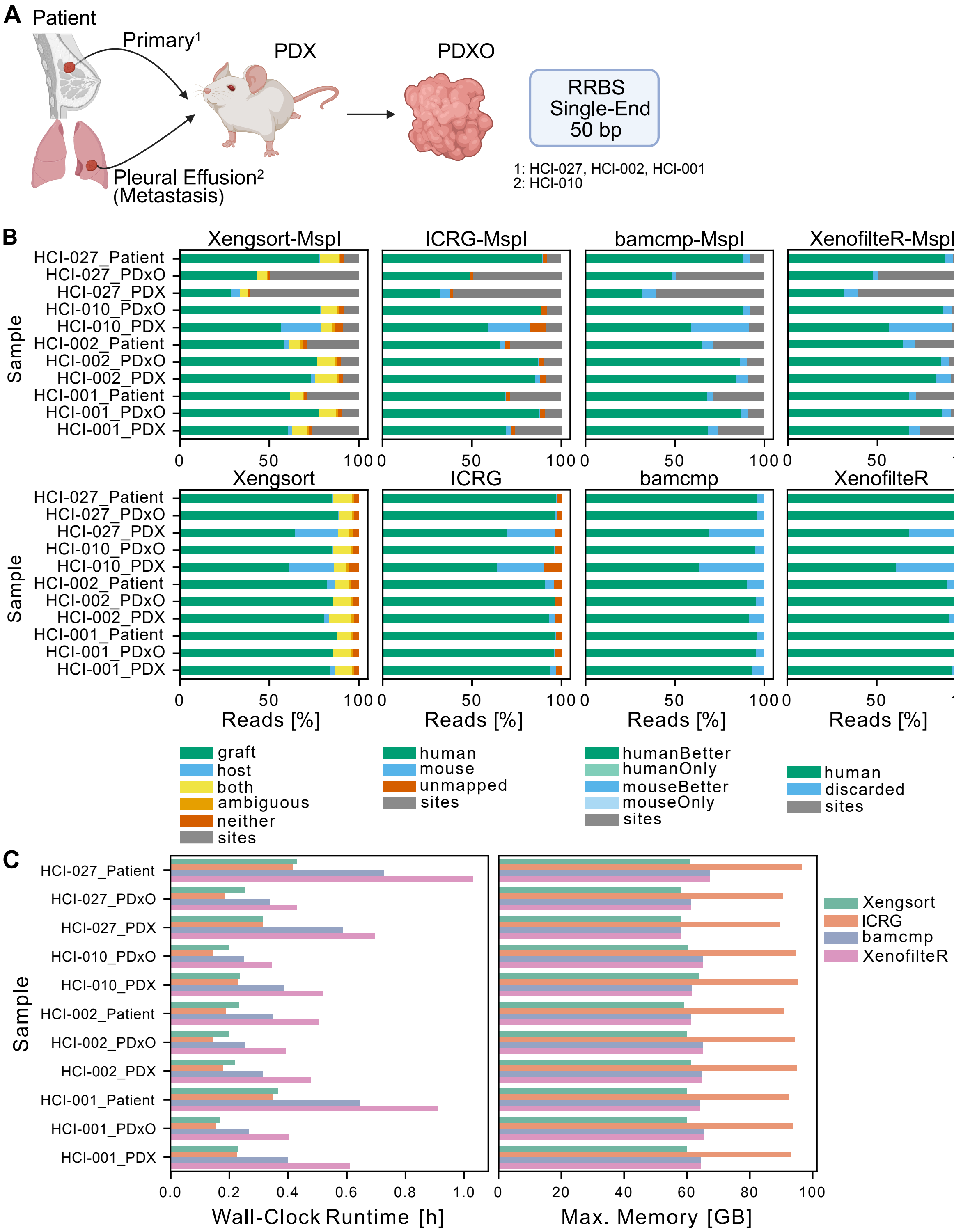


**Extended Data Fig. 4**: Validation of read-sorting algorithms on breast cancer PDX Reduced-Representation Bisulfite-Sequencing (RRBS) data (GSE152202) A. Single-end 50 bp RRBS data included primary and metastatic breast cancer PDXO and PDX samples. B. Classification of sample reads via Xengsort, ICRG, bamcmp, and XenofilteR with MspI start site prefiltering (top) or sorting of all reads (bottom). *sites* (marked in grey) indicate reads not starting with the MspI start site. C. Sum of wall-clock runtime and maximal memory usage for the combined read-sorting and alignment step for each non-MspI-prefiltering workflow.
